# Beyond the (geometric) mean: stochastic models undermine deterministic predictions of bet hedger evolution

**DOI:** 10.1101/2023.07.11.548608

**Authors:** Maya Weissman, Yevgeniy Raynes, Daniel Weinreich

## Abstract

Bet hedging is a ubiquitous strategy for risk reduction in the face of unpredictable environmental change where a lineage lowers its variance in fitness across environments at the expense of also lowering its arithmetic mean fitness. Classically, the benefit of bet hedging has been quantified using geometric mean fitness (GMF); bet hedging is expected to evolve if and only if it has a higher GMF than the wild-type. We build upon previous research on the effect of incorporating stochasticity in phenotypic distribution, environment, and reproduction to investigate the extent to which these sources of stochasticity will impact the evolution of real-world bet hedging traits. We utilize both individual-based simulations and Markov chain numerics to demonstrate that modeling stochasticity can alter the sign of selection for the bet hedger compared to deterministic predictions. We find that bet hedging can be deleterious at small population sizes and beneficial at larger population sizes. This non-monotonic dependence of the sign of selection on population size, known as sign inversion, exists across parameter space for both conservative and diversified bet hedgers. We apply our model to published data of bet hedging strategies to show that sign inversion exists for biologically relevant parameters in two study systems: *Papaver dubium*, an annual poppy with variable germination phenology, and *Salmonella typhimurium*, a pathogenic bacteria that exhibits antibiotic persistence. Taken together, our results suggest that GMF is not enough to predict when bet hedging is adaptive.

## Introduction

When environmental change is unpredictable, the most optimal adaptation can be difficult to predict. Is it better for a lineage to adopt a high risk and high reward strategy by producing offspring that maximize reproductive success in some environments but fail in others? Or to “hedge one’s bets”: either by producing offspring with a phenotype that is a “sure bet” in any environment, or by spreading risk via phenotypically diverse offspring? Due to the multiplicative nature of reproduction, adaptations that reduce risk, such as bet hedging, are especially important (Box 1) ([1], [2], [3], [4]).

Bet hedging is a ubiquitous strategy for risk reduction in varying environments wherein a lineage sacrifices its short term fitness in order to insulate itself against uncertain environmental conditions ([2], [3], [5]). Mathematically, bet hedging is defined as any strategy that lowers fitness variance at the expense of mean fitness (Box 1) ([3]). More specifically, bet hedging increases geometric mean fitness (GMF) at the expense of arithmetic mean fitness (AMF) ([3], [4]). Other possible risk reduction strategies include adaptive tracking and phenotypic plasticity ([6], [5]). Adaptive tracking is the evolution of optimal trait values via standing genetic variation, while phenotypic plasticity is a strategy where individuals change their phenotype based on environmental cues ([7], [5]). Bet hedging is thought to be the most adaptive of these strategies when environmental change is rapid and unpredictable ([5], [8]).

Bet hedging has evolved in over 29 classes across 16 phyla ([5]). The canonical example of bet hedging is delayed seed germination in annual plants, wherein some proportion of seeds enter a seed-bank instead of germinating immediately. Delayed seed germination decreases the average number of seeds that germinate each year, but also lowers the variance in fitness over time by maintaining the seed-bank ([4], [9], [10]). Another well-known example of bet hedging is bacterial persistence, a strategy that allows bacteria to survive antibiotic pressure by producing a subpopulation of persister cells ([11]). Persister cells divide more slowly, thereby reducing the population’s mean fitness, but are able to survive longer periods of antibiotic pressure, increasing their fitness in the presence of antibiotics and thus lowering fitness variance across the two environmental conditions ([12], [13]). Other examples of bet hedging include variable germination phenology in plants ([14], variable egg hatching in crustaceans ([15]), mouthbrooding in cichlids ([16]), altruism in birds ([17]), amphicarpic seed production in annual plants ([18]), and diauxic shift in yeast ([19]).

### Box 1

**Defining types of means and variances**

Different kinds of both mean and variance are important in understanding bet hedging. When the fitness of an allele is not constant over time, determining its long-term evolutionary outcome requires consideration of average fitness. Average fitness over time can be calculated either using the arithmetic mean (AMF) or the geometric mean (GMF). AMF is computed as the sum of a series, divided by the number of elements in the series (*n*). GMF, on the other hand, is computed as the *n*^*th*^ root of the product of the series ([20]). GMF is often used in predicting long term evolutionary success because it better captures the multiplicative nature of reproduction, and therefore the importance of extremely low fitness values and temporal fitness variance ([3]). To visualize the utility of GMF, imagine a series of fitness values that includes a zero at one generation. Biologically, we know that if a lineage has a fitness of zero in one generation, it will go extinct. The arithmetic mean of a series that includes a zero will be low, but not zero. However the geometric mean of this series will be zero, better reflecting the eventual fate of the lineage ([21], [6]).

Variance can either be measured within a generation (i.e. spatial variance) or between generations (i.e. temporal variance). While the effects of both within- and between-generation variance on bet hedging have been studied ([22], most frequently, the benefit of bet hedging is thought of as specifically reducing temporal variance ([2], [9]). This is because temporal fitness variance will have the largest impact on multiplicative reproduction, and therefore, on the long term success of a lineage ([6], ([23]). However, within-generation fitness variance still plays a role in bet hedger evolution, specifically for diversified bet hedgers, which will by definition increase their within-generation fitness variance through the production of multiple phenotypes. While we will discuss both within- and between-generation fitness variance, throughout this paper we focus specifically on environments that only vary in time.

All bet hedging strategies can be classified into one of two categories: conservative or diversified bet hedging ([2], [3], [10], [5]). To illustrate how these two strategies reduce risk, consider a simplified environment that changes unpredictably between a stress-free, nutrient rich state (Environment A) and a high-stress, nutrient-poor state (Environment B) (fig 1A). By definition, the wild-type always expresses a specialist phenotype, which has a high fitness in the stress-free environment, but a low fitness in the high-stress environment. As a result, the wild-type specialist has a high temporal fitness variance (Box 1). The conservative bet hedger reduces variance by having all offspring adopt the same generalist phenotype, which has an intermediate fitness regardless of environment. This means that the conservative phenotype has a lower fitness than the wild-type specialist in the stress-free environment, but a higher fitness than the wild-type in the high-stress environment. Conservative bet hedging strategies therefore lower both the between-generation fitness variance and maintain a low within-generation fitness variance ([3], [5], [23]). On the other hand, the diversified bet hedger reduces risk by producing a range of phenotypically distinct offspring, such that some offspring are better suited to each possible environment. In this way, diversified bet hedgers embody the adage “don’t put all of your eggs in one basket.” Like the conservative bet hedger, the diversified bet hedger has a lower fitness than the wild-type in the stress-free environment, as some proportion of offspring are maladapted, and a higher fitness than the wild-type in the high-stress environment, as some proportion of offspring are well adapted. Unlike the conservative bet hedger, the diversified bet hedger has a non-zero within-generation fitness variance. However, because some proportion of offspring are always better adapted to any environment, the between-generation fitness variance of the diversified bet hedger is lowered ([3], [5], [23]).

**Figure 1:**
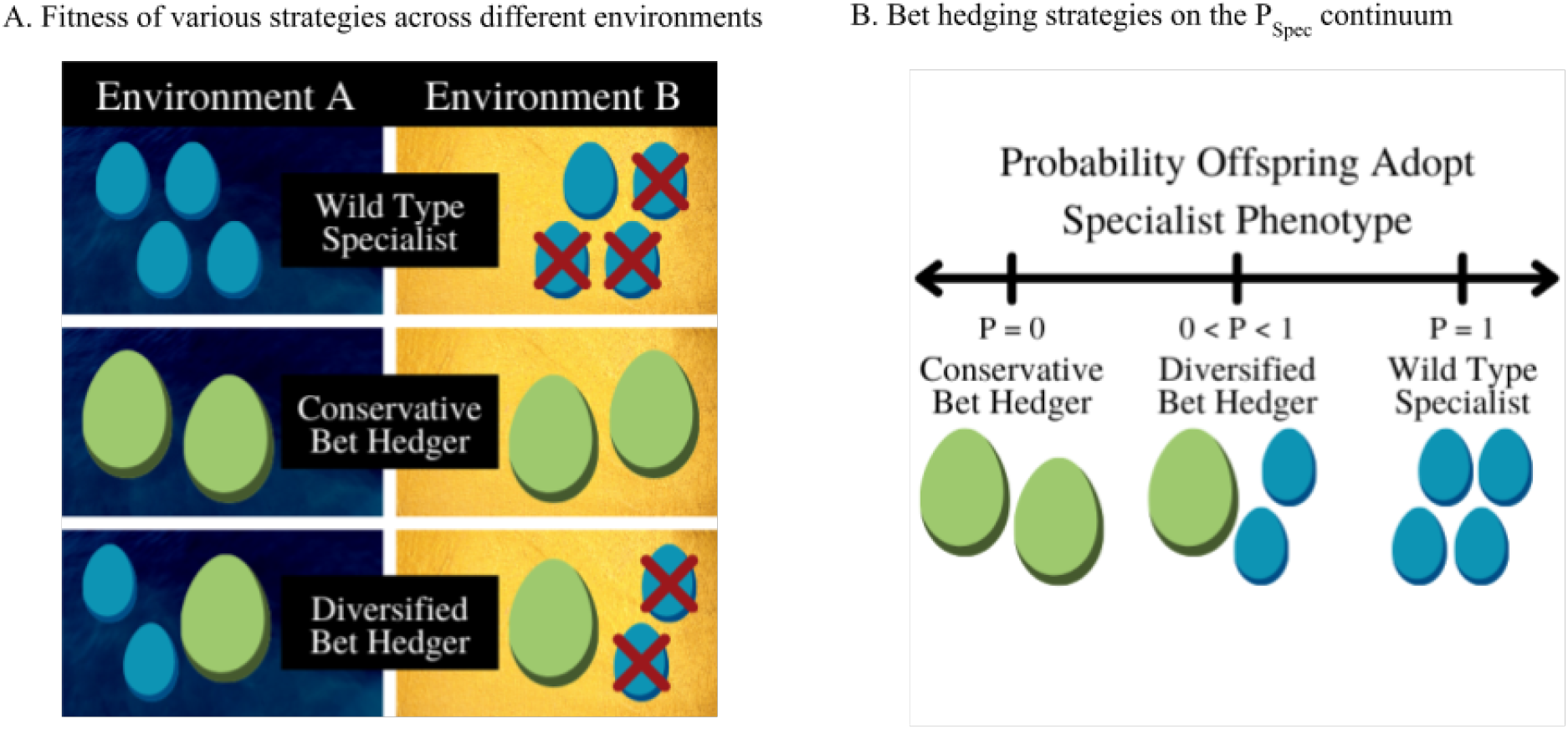
The benefit of bet hedging is seen through the fitness effects of different phenotypes across environmental conditions. A. The environment randomly oscillates between two states: A (stress-free) and B (high-stress). The wild-type specialist has a high fitness in Environment A and a low fitness in Environment B, resulting in both a high AMF and a high fitness variance. The conservative bet hedger produces offspring that all adopt the generalist phenotype, which has an intermediate fitness across both environments. The diversified bet hedger produces phenotypically variable offspring, such that some proportion of offspring are well adapted to the environment. B. All three strategies can exist on a continuum of the proportion of offspring that adopt the specialist phenotype as opposed to the generalist phenotype, or *P*_*Spec*_. For a wild-type lineage, *P*_*Spec*_ = 1. A conservative bet hedging lineage has a *P*_*Spec*_ = 0. By definition, the diversified bet hedger has 0 ¡ *P*_*Spec*_ ¡ 1, where *P*_*Spec*_ proportion of offspring will adopt the specialist phenotype and 1 *− P*_*Spec*_ proportion of offspring will adopt the generalist phenotype.

Diversified bet hedging strategies can be further subdivided into two categories: “high risk /low risk” and “high risk / high risk” ([24]). Returning to our previous example, the diversified bet hedger is a “high risk / low risk” strategy, as one of its two possible phenotypes is the “high risk” specialist phenotype, while the other is the “low risk” conservative phenotype. “High risk / low risk” diversified bet hedging and conservative bet hedging strategies can be regarded as existing along a continuum of possible distributions of phenotypes among offspring ([25], [24], [23]). Many real-world examples of bet hedging strategies fit the “high risk / low risk” model, such as bacterial persistence ([12]), which are often classified as “high risk / low risk” ([24]). However, not all putative bet hedging traits align with this continuum; for some diversified bet hedgers there is no associated conservative strategy possible. These diversified bet hedging strategies are considered “high risk / high risk”, as each phenotype can be considered a specialist in a different environment ([24]). For example, in the delayed seed germination in annual plants there is no assured “generalist” year in which the environment is guaranteed to be good.

Previously, the evolution of bet hedging has largely been explored in deterministic models ([1], [26], [23], [27], [28]). In this deterministic framing, bet hedging is beneficial only when the bet hedger has a greater GMF than the specialist wild-type ([10], [5]). Of note, the GMF is equivalent to assuming that the predictions for reproductive outcome, phenotypic distribution, and environmental regime are precisely realized. However, these assumptions will only be met as population size and time to fixation / loss approach infinity. In reality, since population size and time to fixation / loss are finite, populations will experience variance in stochasticity in reproduction, realized phenotypic distribution, and environment, which can have profound impacts on the evolution of bet hedging.

Stochastic models of bet hedging have begun to explore how this discrepancy between predicted GMF and realized phenotypic distribution, environment, and reproductive output can impact the evolution of bet hedging ([25], [29], [6], [30], [31], [32], [33]). Previous research has found that stochasticity in phenotype can cause bet hedging to be less beneficial than expected by increasing the extinction risk ([33]). Additionally, some studies have suggested that the sign of selection of bet hedging can change with population size ([1], [29], [33]), a phenomenon known as sign inversion ([34], [35]). However, more work is needed to connect stochastic models, deterministic theory (i.e. GMF), and empirical data to predict the adaptive significance of real-world bet hedging traits.

Here, we utilize both individual-based simulations and Markov chain numerics to study the effects of stochasticity on the adaptiveness of bet hedging. We then extend our model across parameter space and show that this phenomenon is expected to impact the evolution of a wide range of conservative, “high risk / low risk” diversified, and “high risk / high risk” diversified bet hedgers. Finally, we apply our model to two species in nature and show that, for biologically realistic parameter values, modeling stochasticity alters the sign of selection for the bet hedger relative to GMF based predictions. We find that both systems exhibit sign inversion, or are beneficial at large population sizes and deleterious at small population sizes. As the largest existing issue in the field is determining when putative bet hedging strategies are adaptive, we propose that using GMF alone is not sufficient to predict when a trait is beneficial.

## Methods

### Code availability

All code is available at https://github.com/mweissman97/bethedging_stochastic.

### Stochastic individual based simulations

We model the evolution of bet hedging in asexual populations of constant size *N* evolving in discrete non-overlapping generations under the Wright-Fisher model. Simulations start with *N* individuals, consisting of a single invading bet hedging mutant and *N −* 1 resident wild-type individuals, and end when the bet hedger reaches fixation (frequency of 100%) or goes extinct (frequency of 0%). Every generation, the environment adopts one of two states: *EnvironmentA*, or the stress-free condition, with probability *P*_*A*_ and *EnvironmentB*, or the stress condition, with probability 1 *− P*_*A*_. We keep *P*_*A*_ = 0.5 constant, as our results are not sensitive to the value of *P*_*A*_. The wild-type phenotype has a higher fitness in *A* (*w*_*A*_ = 2), and a much lower fitness in *B* (*w*_*B*_ = 0.5). The GMF of the wild-type is thus 2^0.5^ *×* 0.5^0.5^ = 1.

In the “high risk / low risk” model of diversified bet hedging, the fitness of the bet hedger is defined by two parameters: *w*_*C*_, the fitness of the conservative phenotype in both environments, and *P*_*Spec*_, the proportion of offspring that adopt the specialist phenotype. When *P*_*Spec*_ = 0, all offspring will adopt the conservative phenotype, and the lineage will exhibit a conservative bet hedging strategy. When *P*_*Spec*_ = 1, the lineage will exhibit the wild-type specialist strategy. For intermediate values of *P*_*Spec*_, the lineage will exhibit a diversified bet hedging strategy. From these two parameters, we can calculate the GMF of the bet hedger as:

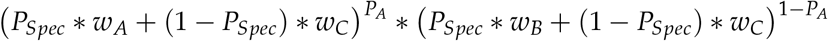

For a conservative bet hedger, since *P*_*Spec*_ = 0, the GMF will be simply *w*_*C*_. Parameters are selected such that the bet hedger always has both a lower AMF and temporal fitness variance than the resident wild-type. We present the GMF of the bet hedger as Δ*GMF*, or the GMF of the bet hedger minus the GMF of the wild-type.

When the diversified bet hedger utilizes a “high risk / high risk” strategy, the second phenotype is a B environment specialist, as opposed to the conservative phenotype. As there are now two different specialist phenotypes, we will refer to the previously described environment A specialist as the “*α* specialist” and the environment B specialist as the “*β* specialist”. The *β* specialist has a high fitness in environment B (*w*_*β,B*_), and a low fitness in environment A (*w*_*β,A*_). The GMF of the “high risk / high risk” diversified bet hedger is thus:

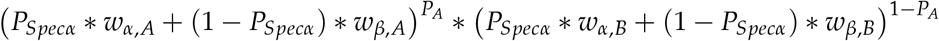

In order to precisely control the Δ*GMF* of the “high risk / high risk” diversified bet hedger, the fitness values of the *β* specialist in both environments are calculated as follows. First, the fitness of the *β* specialist in environment B is assigned as *w*_*β,B*_ = 2, equal to the maximum possible fitness of the *α* specialist. The *β* specialist’s fitness in environment A (*w*_*β,A*_) is then calculated to yield the desired Δ*GMF*, given *P*_*Specα*_ and the other fitness values. In cases where the calculated value of *w*_*β,A*_ *<* 0, *w*_*βA*_ is set to 0.01, and the appropriate value of *w*_*βB*_ is then calculated.

Reproduction is simulated according to the Wright–Fisher model, where the number of offspring in the next generation is drawn from a multinomial distribution with the expectation determined by each lineage’s current frequency multiplied by the lineage’s relative fitness in that generation. The probability of fixation is calculated as the proportion of replicates in which the bet hedger reaches fixation. We follow others ([36], [37], [34]) and normalize the probability of fixation by the expected probability of fixation for a neutral mutation. As all simulations begin with a single bet hedger, the normalized probability of fixation is algebraically *P*_*Fix*_/(1/*N*).

Individual-based simulations are carried out in the Julia Programming Language ([38]).

#### Implementing phenotypic stochasticity in individual based simulations

Phenotypic stochasticity can either be turned on or turned off in our simulations. When pheno-typic stochasticity is turned on, every generation the realized number of offspring that adopt the specialist phenotype is drawn from a binomial distribution with probability equal to *P*_*Spec*_ and sample size equal to the number of bet hedger individuals in the population.

When phenotypic stochasticity is turned off, every generation the fitness of all bet hedger individuals is equal to the expected fitness given *P*_*Spec*_. Regardless of bet hedger counts, *P*_*Spec*_ is always precisely equal to its expectation, even if a fraction of an individual would need to adopt the specialist phenotype in order to do so. This essentially renders the bet hedger a conservative bet hedger, with some variance in fitness between the two environments.

#### Implementing environmental stochasticity in individual based simulations

Environmental stochasticity can also either be turned on or off in our simulations. When environmental stochasticity is turned on, every generation the environmental state is randomly assigned to either Environment A (with probability *P*_*A*_), or Environment B (with probability 1 *− P*_*A*_).

When environmental stochasticity is turned off, the environment switches deterministically every generation. In the first generation, the environment is randomly assigned to either Environment A (with probability *P*_*A*_), or Environment B (with probability 1 *− P*_*A*_). As we keep *P*_*A*_ = 0.5 fixed in our simulations, the environment in subsequent generations is assigned as opposite to the environment in the previous generation.

#### Markov matrix numeric model

We also use a Markov numeric matrix model to validate our simulations by again computing the probability of fixation for a single bet hedger in a population of *N −* 1 wild-type individuals.

First, single generation transition matrices are constructed for each possible environment (designated as *T*_*A*_ and *T*_*B*_, for environments A and B respectively). Transition matrices are *N* +1 *× N* + 1 matrices that give the probability of transitioning from all possible values of starting count, *x*_*t*_, to all possible values of ending count *x*_*t*+1_ in a single generation, given the fitness of both the wild-type and the mutant. For more information on constructing transition matrices, see the supplement.

Because GMF is a measure of fitness across environments, and therefore generations, it cannot be realized in any single generation. As such, we introduce an effective generation matrix. We define an effective generation as the minimum number of generations necessary for all environments to be experienced with their expected proportion (*P*_*A*_). In our case, as *P*_*A*_ = 0.5, an effective generation is two consecutive generations. The effective generation regime matrices are therefore calculated as the product of the transition matrices for the first and second generations (*T*_*EnvinGeneration*=2_ *× T*_*EnvinGeneration*=1_). This yields a matrix that will give the expected transition probabilities at the end of the effective generation. Effective generation regime matrices are computed separately for each possible environmental regime (*AB, BA, AA*, and *BB*), and then averaged together.

Finally, the probability of fixation is calculated by estimating the stationary distribution of the effective generation matrix. The stationary distribution (*π*) is the matrix that satisfies the condition *π* = *π × T*. The probability of fixation is the *P*(*x*_*t*_ = 1|*x*_*t*+1_ = *N*) entry of the stationary distribution matrix.

Matrix multiplication was carried out using the MATLAB Programming Language ([39]).

#### Implementing phenotypic stochasticity in the Markov matrix model

Phenotypic stochasticity can change the realized fitness of the bet hedger in any single generation. When phenotypic stochasticity is turned on, the realized phenotypic distribution of the bet hedger can differ from the expectation. As variance is modulated by the number of bet hedgers present, or *x*_*t*_, we must calculate all possible values of realized *P*_*Spec*_ for every column in our matrix individually, using a binomial distribution with probability equal to *P*_*Spec*_ and sample size equal to *x*_*t*_. For each possible value of realized *P*_*Spec*_, we then calculate 1) the probability of realizing said *P*_*Spec*_, and 2) all transition probabilities of reaching a count of *x*_*t*+1_ in the next time step, given the fitness of the bet hedger associated with the realized *P*_*Spec*_. We then take the weighted average of all possible transition probabilities for the *x*_*t*_ column. We repeat this for all columns, until we have a full transition matrix for both environmental conditions.

When phenotypic stochasticity is turned off, the realized phenotypic distribution of the bet hedger is precisely equal to the expectation (*P*_*Spec*_). Thus, the fitness of the bet hedger is assumed to be constant, and independent of bet hedger count, for each of the environments.

#### Implementing environmental stochasticity in the Markov matrix model

Environmental stochasticity changes the exact sequence of environments experienced by the bet hedger, and therefore the environmental regimes possible within a single effective generation. When environmental stochasticity is turned “on”, there are four possible environmental regimes for the effective generation: *AB, BA, AA*, or *BB*. The effective generation matrix is therefore the arithmetic mean of the *T*_*B*_ *× T*_*A*_, *T*_*A*_ *× T*_*B*_, *T*_*A*_ *× T*_*A*_, and *T*_*B*_ *× T*_*B*_ regime matrices.

When environmental stochasticity is turned off, as the environment switches every generation, there are only two possible environmental regimes for the effective generation: *AB* or *BA*. It follows that the effective generation matrix will be the arithmetic mean of the *T*_*B*_ *× T*_*A*_ and *T*_*A*_ *× T*_*B*_ regime matrices.

### Constructing parameter space surveys

We utilize our stochastic, individual-based simulations to construct large scale parameter space surveys, since their run times scale more favorably as population size increases. For every point in parameter space, simulations were carried out at 20 different populations sizes spaced evenly on a logarithmic scale between 10^0^ and 10^6^ to construct a normalized probability of fixation vector.

We vary both the value of *P*_*Spec*_ from 0 to 1, and the value of Δ*GMF* from -0.15 to 0.125. In order to vary Δ*GMF*, we calculate the corresponding fitness values of the second phenotype, either *w*_*C*_ for the “high risk / low risk” model or [*w*_*βA*_, *w*_*βB*_] for the “high risk / high risk” model, that will yield the appropriate Δ*GMF* given the value of *P*_*Spec*_. However, for certain values of *P*_*Spec*_, there is no possible value of *w*_*C*_ that will yield certain values of Δ*GMF*. For example, when *P*_*Spec*_ = 1.0, no individuals will adopt the conservative phenotype, and therefore it is impossible to have a value of Δ*GMF/*= 0. For combinations of parameter values that are impossible to experience, the point in parameter space is intentionally left blank.

#### Characterizing the sign of selection of probability of fixation vectors

After calculating the normalized probability of fixation (*NP*_*Fix*_) as a function of population size to a maximum population size of 10^6^, the vector of all *NP*_*Fix*_ values is classified based on whether the sign of selection is unconditionally beneficial, unconditionally deleterious, neutral, or exhibits sign inversion computationally as follows. First, at each population size, the probability of fixation is evaluated as being significantly different from the neutral expectation using a *p <* 0.01 confidence interval. The confidence interval is found by constructing a binomial probability distribution with probability equal to the probability of fixation for a neutral mutation and a number of trials equal to the number of replicates. Then, each *NP*_*Fix*_ vector is evaluated to assay the sign of selection. A vector is labeled neutral if all points are not significantly different from the neutral expectation. A vector is classified as beneficial if all points on the vector are greater than or equal to 1, or deleterious if all points are less than or equal to 1. A vector is labeled as exhibiting sign inversion if a) when *N >* 1, the sign of selection changes only once (i.e. from deleterious to beneficial, or from beneficial to deleterious), b) if the sign of the slope of the vector changes only once. *NP*_*Fix*_ vectors that do not meet any of these criteria were plotted and evaluated visually.

#### Interpolating the value of N_Crit_

For curves that exhibit sign inversion, we interpolate the value of *N*_*Crit*_, or the critical population size where the probability of fixation crosses the neutral expectation, using the Linear Interpolation function from the Interpolations Library in the Julia Programming Language ([38].

### Empirical data parameter estimates

Fitness and life history measurements for *Papaver dubium* were taken from Arthur 1973 ([14]). Parameters for *Salmonella typhimurium* were taken from Arnoldini 2014 ([40]). Fitness values were normalized such that the fitness of the conservative phenotype in both environments was 1 (*w*_*C*_).

A range of effective population size for both species was estimated to the nearest order of magnitude. The population size of *Papaver dubium* was estimated using a combination of the census size of the population surveyed ([14]) and estimates of *N*_*e*_/*N* from Heywood 1986 ([41]). The effective population size (*N*_*e*_) of *S. typhimurium* was assumed to be comparable to other free living bacteria ([42]).

The parameter estimates used are summarized in the table below:

| Species | $P_{Spec}$ | $w_A$ | $w_B$ | $N_e$ range |
| --- | --- | --- | --- | --- |
| *Papaver dubium* | 0.53 | 18 | 0.5 | $10^2 - 10^4$ |
| *Salmonella typhimurium* | 0.81 | 1.8 | 0.075 | $> 10^8$ |

#### Climate estimates for P. dubium

Life history parameters for *P. dubium* were measured from 1966 - 1970 in Birmingham, England ([14]). As such, we utilize daily mean temperature measurements for Central England to estimate the probability of a mild winter ([43]). We constructed distributions of the mean winter temperature, the number of days below 0°C per winter, and the number of days below -1.6°C per winter (supplemental figure 1). In Arthur 1973 ([14]), it was established that 1966 and 1967 were considered “mild years”, while 1968 was a “harsh year”. We estimated the threshold between harsh and mild years as being the mean of 1968 and 1966 (the cooler of the two mild years) measurements. The proportion of mild winters was then estimated as the proportion of years with higher mean temperature, less days below 0°C, or less days below -1.6°C compared to the threshold.

### Statistical analyses

Statistical analyses were carried out using the R Programming Language ([44]).

## Results

### Stochasticity changes the extent to which bet hedging is beneficial

Under a purely deterministic population genetics model, beneficial traits will always reach fixation and deleterious traits will always be lost. In this framing, whether or not the bet hedger is adaptive is determined by the GMF of both strategies: the strategy with the higher GMF will always reach fixation ([3]). However in a stochastic model, we instead consider the probability of fixation for an invading mutant to assay its sign of selection. Following others ([36], [37], [34]), we normalize the probability of fixation by the neutral expectation so that the neutral benchmark is always equal to 1. As the starting frequency of the bet hedger is always 1/*N*, the normalized probability of fixation is *P*_*Fix*_/(1/*N*). A beneficial trait will have a normalized probability of fixation greater than the neutral expectation (*NP*_*f ix*_ *>* 1), while a deleterious trait will have a normalized probability of fixation less than the neutral expectation (*NP*_*f ix*_ *<* 1).

We utilize a probabilistic model that incorporates three sources of stochasticity: stochasticity in reproduction, phenotypic distribution, and environment. Classical population genetics theory has established the role of stochasticity in reproduction, called genetic drift. As population size increases, the effect of genetic drift decreases and the efficiency of selection increases ([45]). Stochasticity in phenotype is randomness in the distribution of phenotypes amongst diversified bet hedger offspring within a generation. By definition, a conservative bet hedger does not experience any stochasticity in phenotype, as all conservative offspring will adopt the same, low risk phenotype. Previous research has determined explicitly incorporating phenotypic stochasticity can increase the extinction risk for bet hedging ([33]). Finally, stochasticity in the environment is randomness in the sequence of environments experienced by a lineage. It has also been shown that environmental stochasticity can cause the extinction risk for bet hedging to decrease, but only at small population sizes ([33]). But the question remains: are these sources of stochasticity enough to impact the real-world evolution of bet hedging traits, or is GMF sufficient to predict when bet hedging traits are expected to evolve?

We first focus on the “high risk / low risk” model of diversified bet hedging.

We find the sign of selection for a single invading diversified bet hedger mutant (*P*_*Spec*_ = 0.5) in a population of *N −* 1 wild-type specialist individuals (*P*_*Spec*_ = 0, *GMF* = 1) (fig 2). The fitness of the bet hedger is characterized by Δ*GMF*, or the GMF of the bet hedger subtracted from the GMF of the wild-type. Therefore, when Δ*GMF >* 0, the bet hedger has a greater GMF and on deterministic considerations, is predicted to be beneficial. Representative values of Δ*GMF* are selected to show the probability of fixation when bet hedging is predicted to be strongly beneficial (Δ*GMF* = 0.1), weakly beneficial (Δ*GMF* = 0.01), neutral (Δ*GMF* = 0), weakly deleterious (Δ*GMF* = -0.015), and strongly deleterious (Δ*GMF* = -0.1). We utilize both stochastic individual-based simulations and Markov matrix numerics, and find no difference between the two model types in results (*p* = 0.9964).

**Figure 2:**
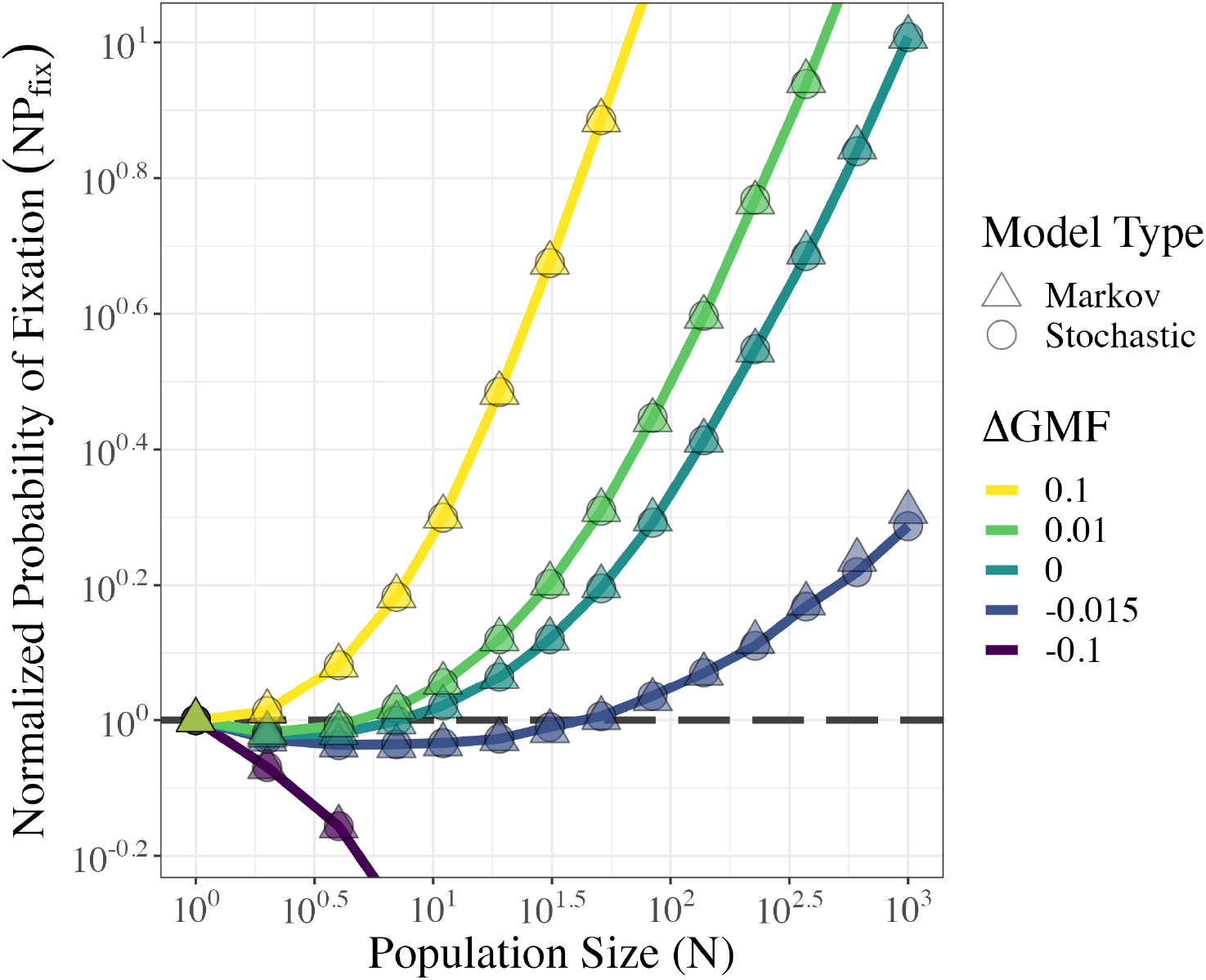
The normalized probability of fixation (*NP*_*Fix*_) for a single invading “high risk / low risk” diversified bet hedging mutant (*P*_*Spec*_ = 0.5) in a population of *N*− 1 wild-type individuals (*P*_*Spec*_ = 1.0) is shown as a function of population size and Δ*GMF*. We find no significant difference between the stochastic, individual-based simulations (circles) and Markov matrix numerics (triangles) (*p >* 0.99). When Δ*GMF* = 0.1 and -0.1, our results agree with the deterministic prediction that bet hedging is beneficial (*NP*_*f ix*_ *>* 1) or deleterious (*NP*_*f ix*_ *<* 1) across all population sizes examined. For the three intermediate values of Δ*GMF* (0.01, 0, and -0.015), we find that the sign of selection disagrees with deterministic predictions. Instead, bet hedging is deleterious at small population sizes (*NP*_*f ix*_ *<* 1) and beneficial at larger population sizes (*NP*_*f ix*_ *>* 1). It thus exhibits sign inversion.

When bet hedging is expected to be strongly beneficial (here, Δ*GMF* = +0.1), we find that bet hedging is unconditionally beneficial (*NP*_*f ix*_ *≥* 1), which aligns with the deterministic prediction. Similarly, when bet hedging is expected to be strongly deleterious (here, Δ*GMF* = *−*0.1), the sign of selection also agrees with the deterministic prediction and bet hedging is deleterious across all population sizes (*NP*_*f ix*_ *≤* 1).

When the GMF of the bet hedger is equal to that of the resident wild-type (Δ*GMF* = 0), bet hedging should be neutral according to the deterministic prediction. Instead, we find that the normalized probability of fixation is below the neutral expectation at small population sizes, crosses the neutral expectation (*N ≈* 10), and then becomes greater than the neutral expectation at larger population sizes. More simply, when Δ*GMF* = 0, bet hedging is deleterious at small population sizes and beneficial at larger population sizes. This non-monotonic dependence of the sign of selection on population size is known as sign inversion ([34]). We also find that when the bet hedger is weakly beneficial or weakly deleterious (Δ*GMF* = +0.01 and Δ*GMF* = *−*0.015 respectively), bet hedging continues to exhibit sign inversion. When Δ*GMF* = 0.01, we expect bet hedging to be unconditionally beneficial, but it is instead deleterious at sufficiently small population sizes. Conversely, when Δ*GMF* = *−*0.015 and bet hedging is expected to be unconditionally deleterious, bet hedging is beneficial at sufficiently large population sizes.

### Sign inversion exists across parameter space

We extend our generalized model across parameter space to determine the extent to which the sign of selection for the bet hedger differs from the deterministic prediction. We characterize the sign of selection across *P*_*Spec*_ on the horizontal axis (which captures the continuum of conservative, diversified, and specialist strategies) and Δ*GMF* on the vertical axis (ranging from expected to be deleterious to beneficial). We maintain *P*_*A*_ = 0.5. Under a “high risk / low risk” model: *P*_*Spec*_ = 0 corresponds to a conservative strategy, *P*_*Spec*_ = 1 to the wild-type specialist (and is thus neutral), and intermediate values of *P*_*Spec*_ to a diversified strategy. Δ*GMF* is varied by changing the fitness of the conservative phenotype (*w*_*C*_), with Δ*GMF* = 0 where bet hedging is expected to be neutral denoted with a blue line. Values of Δ*GMF* that are impossible to experience in combination with certain values of *P*_*Spec*_ are purposefully left blank (e.g. when *P*_*Spec*_ = 1, the only possible value of Δ*GMF* = 0). For points in parameter space where bet hedging exhibits sign inversion, we interpolate the population size where the *NP*_*f ix*_ curve crosses the neutral benchmark before becoming beneficial (*N*_*Crit*_). For example, in Figure 2, we observed that the *NP*_*f ix*_ curve for Δ*GMF* = 0 crosses the neutral benchmark at an *N*_*Crit*_ *≈* 10.

In our original model that includes stochasticity in both phenotype and environment (fig. 3A), we find that the sign of selection of the bet hedger differs from the deterministic prediction across parameter space. When Δ*GMF >* 0 and bet hedging is predicted to be unconditionally beneficial, bet hedging can instead be deleterious at sufficiently small population sizes. This decrease in fixation probability at small population sizes is driven by phenotypic stochasticity, in agreement with previous findings (supplemental fig. 2A, [33]). On the other hand, when Δ*GMF <* 0 and bet hedging is predicted to be unconditionally deleterious, bet hedging can instead be beneficial at sufficiently large population sizes. This increase in fixation probability at large population sizes is driven by environmental stochasticity (supplemental fig. 2B). While previous research suggested that environmental stochasticity can elevate the probability of fixation at small population sizes ([33]), we instead show that the probability of fixation increases most at large population sizes. We show that this increase in probability of fixation is widespread; the fitness disadvantage of the bet hedger can be as large as Δ*GMF* = *−*0.1, which is analogous to a 10% selective advantage, and still be beneficial at some population sizes.

**Figure 3:**
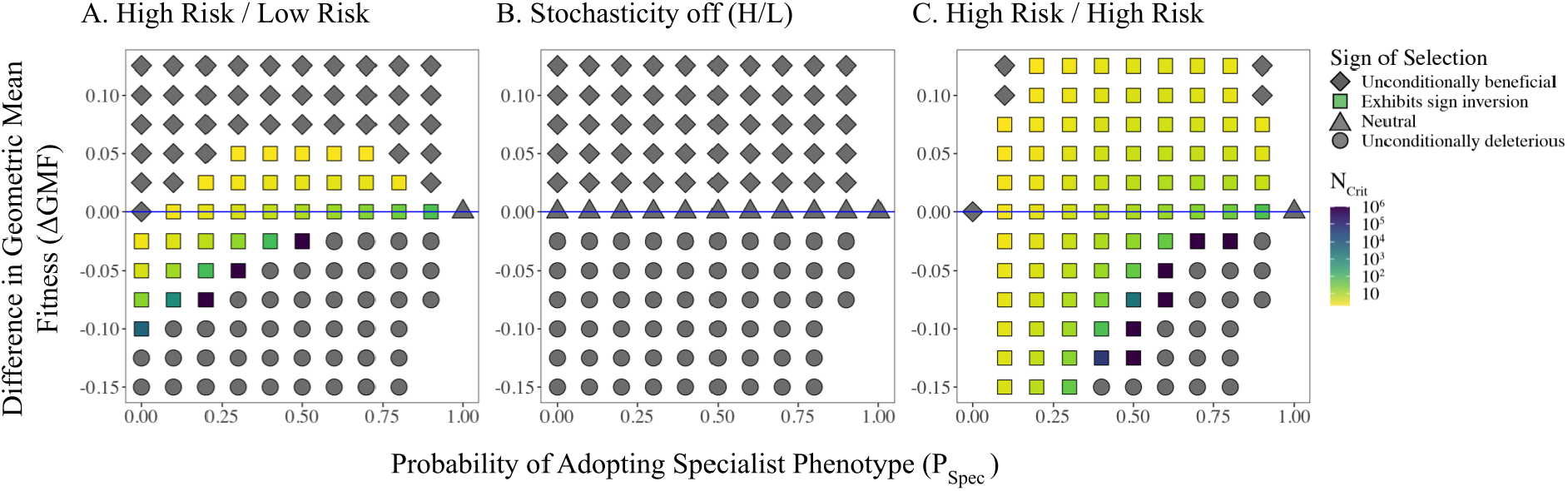
The sign of selection of the bet hedger across parameter space is characterized using stochastic, individual based simulations. The x-axis is *P*_*Spec*_, or the proportion of bet hedger offspring which adopt the specialist phenotype. The y-axis is the difference in geometric mean fitness (Δ*GMF*) of the bet hedger and the wild-type. Values of Δ*GMF* that are impossible to achieve in combination with their associated *P*_*Spec*_ are intentionally left blank (see Methods). For points in parameter space where bet hedging exhibits sign inversion, *N*_*Crit*_ is interpolated. A. In a “high risk / low risk” model of bet hedging, sign inversion occurs across parameter space. Here, *P*_*Spec*_ = 0 corresponds to a conservative bet hedging strategy, 0 *< P*_*Spec*_ *<* 1 corresponds to a diversified bet hedging strategy, and *P*_*Spec*_ = 1 corresponds to the specialist wild-type. B. For the “high risk / low risk” model, when neither stochasticity in environment nor phenotype are incorporated, the sign of selection aligns with the deterministic predictions. C. In a “high risk / high risk” model of diversified bet hedging, we similarly see sign inversion across parameter space. Here, *P*_*Spec*_ = 0 corresponds to a B environment specialist, 0 *< P*_*Spec*_ *<* 1 corresponds to a diversified bet hedging strategy, and *P*_*Spec*_ = 1 corresponds to the A environment specialist wild-type.

When bet hedging exhibits sign inversion, we find that the value of *N*_*Crit*_ increases as the Δ*GMF* decreases. This is unsurprising, as decreasing GMF is expected to cause the probability of fixation to decrease across all population size. This shifting of the *NP*_*f ix*_ curve downward will in turn increase *N*_*Crit*_. When bet hedging has a Δ*GMF >* 0, and phenotypic stochasticity causes bet hedging to be less beneficial than expected, we observe *N*_*Crit*_ values between 2 and 30. Given the narrow range of population sizes where phenotypic stochasticity causes bet hedging to be deleterious, we predict that it is unlikely that phenotypic stochasticity will significantly hinder the evolution of real-world bet hedging strategies. On the other hand, when Δ*GMF <* 0, and environmental stochasticity causes bet hedging to be more beneficial than expected, we observe *N*_*Crit*_ values between 30 and 10^6^. Due to the fact that the greatest observed value of *N*_*Crit*_ is not significantly less than the maximum population size surveyed, it is possible that sign inversion could occur at some points currently labelled as being unconditionally deleterious. However, because *N*_*Crit*_ increases exponentially, extending the maximum population size surveyed to capture this is not feasible (supplemental figure 3). Given the large range of both Δ*GMF* and realistic population sizes where bet hedging is more beneficial than deterministic predictions, we predict that this phenomenon is likely to promote bet hedger evolution in a range of contexts.

When both stochasticity in the phenotype and environment are turned off (fig. 3B), the sign of selection for the bet hedger aligns exactly with the deterministic predictions across all parameter space. We find that bet hedging is unconditionally beneficial when Δ*GMF >* 0, neutral when Δ*GMF* = 0, and unconditionally deleterious when Δ*GMF <* 0. This demonstrates that stochasticity in phenotype and environment must drive the sign of selection for the bet hedger to differ from the deterministic prediction.

We now turn to the “high risk / high risk” model of diversified bet hedging (fig. 3C). In this parameter space, no conservative bet hedging strategy is possible. Therefore, *P*_*Specα*_ = 0 corresponds to a population of *β* B environment specialists, *P*_*Specα*_ = 1 to a population of *α* A environment specialists (and is thus neutral), and intermediate values of *P*_*Spec*_ to a diversified strategy. We continue to vary Δ*GMF* by modulating the fitness of the B Specialist in both environments. Again, for areas of parameter space where the combination of *P*_*Spec*_ = and Δ*GMF* are impossible, points are intentionally left blank. In the “high risk / high risk” model of bet hedging, sign inversion is even more widespread (fig. 3C). We see sign inversion occur both for values of Δ*GMF >* 0.05 and Δ*GMF < −*0.1. This is because “high risk / high risk” bet hedgers will experience greater variance in fitness both within and between generations.

Regardless of the type of bet hedging strategy, conservative vs. diversified or “high risk / low risk” vs. “high risk / high risk”, we show that GMF is not sufficient to predict the sign of selection.

### Stochastic models explain the adaptive significance of real-world bet hedging traits

We parameterize our model using empirical data from two study systems: variable germination phenology in *Papaver dubium* and antibiotic persistence in *Salmonella typhimurium*. These two bet hedging traits were selected because they provided adequate life history measurements, but initial studies were unable to prove the bet hedging trait was adaptive to its environmental regime ([5], [14], [40]).

#### Germination phenology in Papaver dubium

*Papaver dubium*, an annual poppy native to Central England, engages in a type of “high risk / low risk” diversified bet hedging strategy where some proportion of seeds germinate in the fall, while the remainder germinate in the spring ([14]). Fall germinators are akin to our specialist phenotype: they grow much larger and have significantly higher fitness when winters are mild, but have much lower survivorship when winters are harsh. Spring germinators, on the other hand, correspond to our conservative phenotype, as their fitness is largely unaffected by the winter. We utilize empirical measurements for the proportion of seeds that germinate in the fall (*P*_*Spec*_), as well as the fitness of both seed types across two years with mild winters and one year with a harsh winter (*w*_*A*_, *w*_*B*_, and *w*_*C*_) ([14]).

To determine the environmental context in which this strategy evolves, we use historic climate data from the years 1772 - 1973 to estimate the proportion of years with mild winters (or *P*_*A*_ in our model) ([43]). From daily mean temperatures, we analyzed three metrics to estimate *P*_*MildWinter*_: the mean winter temperature, the number of days below 0°C, and the number of days below -1.6°C (which corresponds to a “deep freeze”). While this temperature data is continuous, we utilize a discrete environmental model with only “mild” and “harsh” years. Given the climate observations from Arthur 1973, we estimate the cut-off between our two states to be the midpoint between the observed metrics for 1966 (the less mild winter observed) and 1968 (the one harsh winter observed). From this empirical data, we estimate *P*_*MildWinter*_ to be between 0.42 and 0.59 (supplemental figure 1).

Given these estimates, we find that deterministic estimates would not predict that bet hedging in *P. dubium* is beneficial (fig 4A). We calculate the Δ*GMF* of an invading bet hedging mutant into a population of exclusively fall germinating *P. dubium* across all values of *P*_*MildWinter*_. We find that bet hedging is expected to be deleterious (Δ*GMF <* 0) when *P*_*MildWinter*_ *>* 0.396. Therefore, for the entirety of our range of realized *P*_*MildWinter*_, bet hedging is not expected to evolve according to its GMF.

**Figure 4:**
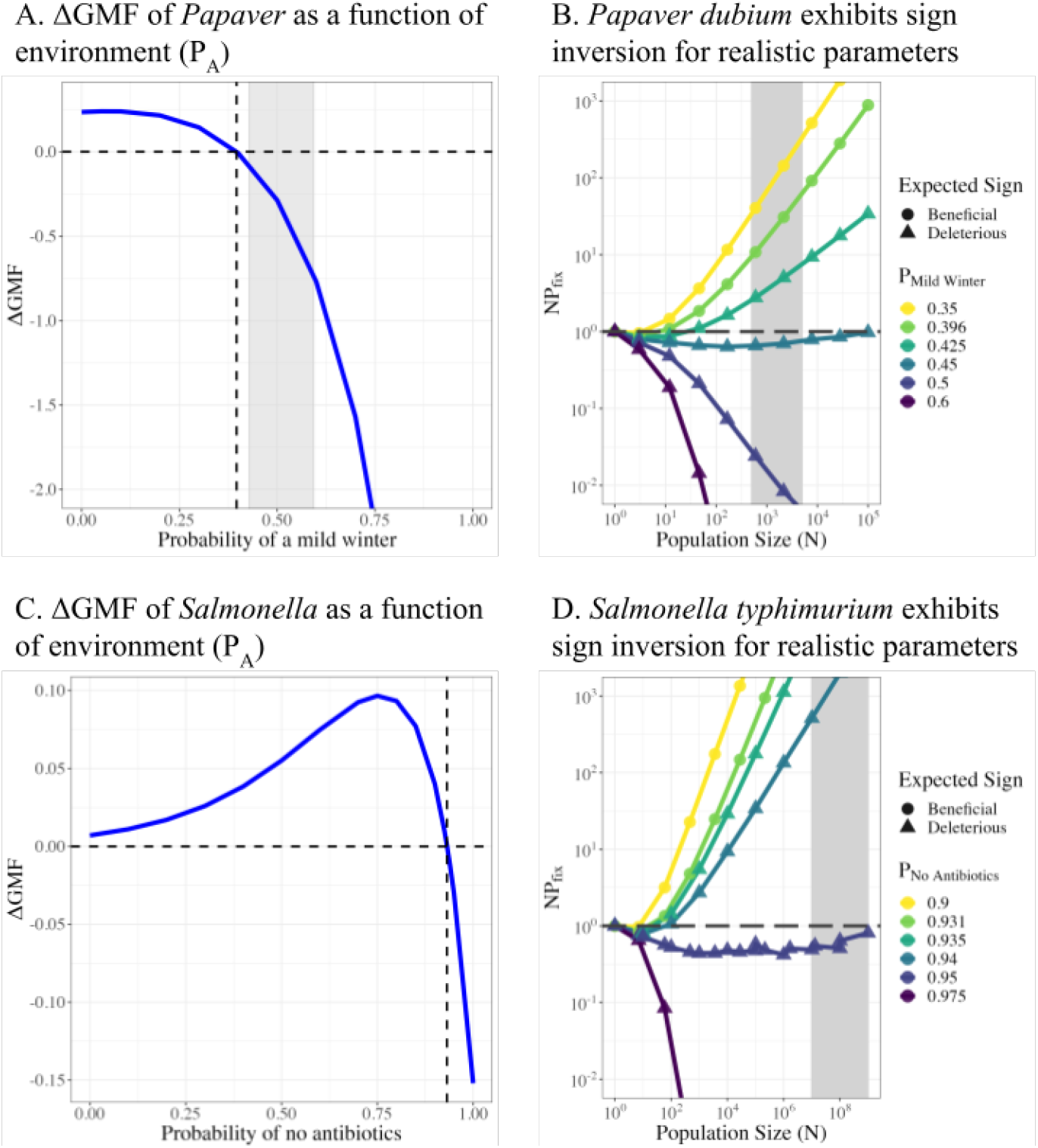
A. Using parameter estimates for bet hedging in *P. dubium* phenology, Δ*GMF*, or the GMF of the bet hedger subtracted by the GMF of a specialist species that exclusively germinates in the fall. Δ*GMF* is caclulated as a function of the proportion of mild winters (*P*_[_ *MildWinter*]). Δ*GMF* predicts bet hedging will be deleterious when *P*_[_ *MildWinter*] *>* 0.396, which includes the entirety of the realistic range of *P*_[_ *MildWinter*] estimated from historical climate data (grey shaded region). B. In contrast, stochastic models predict bet hedging in *P. dubium* will be beneficial within our range of *P*_[_ *MildWinter*] and effective population sizes (grey shaded region). Bet hedging also exhibits sign inversion within this region of parameter space. C. We parametrize our model with life history data from *S. typhimurium* to calculate Δ*GMF*, or the GMF of the bet hedger subtracted by a specialist species with no fraction of persister cells. Δ*GMF* is calculated as a function of the probability of taking no antibiotics (*P*_[_*NoAntibiotics*). Bet hedging is expected to be deleterious when *P*_[_*NoAntibiotics*] *>* 0.932. D. Under a stochastic model, *S. typhimurium* does exhibit sign inversion. With the range of realistic population sizes, bet hedging in *S. typhimurium* is more beneficial than expected in a range of environmental regimes (0.932*P*_[_*NoAntibiotics*] *<* 0.95).

Yet using our stochastic framework, bet hedging in *P. dubium* can evolve (fig 4B). We find that bet hedging is expected to be beneficial for values of *P*_*MildWinter*_ *<* 0.45 within our range of expected population sizes. Additionally, we find that bet hedging exhibits sign inversion, or is deleterious at small population sizes and beneficial at larger population sizes. *P. dubium* exhibits sign inversion both when Δ*GMF <* 0 and when Δ*GMF >* 0. When *P*_*MildWinter*_ = 0.425, and bet hedging is expected to be unconditionally deleterious, our results show that bet hedging can be beneficial when *N >* 50. As this *N*_*Crit*_ is well below the effective population size of *P. dubium*, our model predicts bet hedging to be beneficial in this region of parameter space.

When *P*_*MildWinter*_ = 0.396, and bet hedging is expected to be beneficial across all population sizes, instead, bet hedging can be deleterious when *N <* 10. While estimates of *P. dubium* effective population size are far larger, our results confirm findings that phenotypic stochasticity can cause bet hedging to be less beneficial than expected ([33]). Our results indicate that incorporating stochasticity in environment and phenotype are necessary to explain the evolution of variance in germination timing in *P. dubium*.

### Antibiotic persistence in Salmonella typhimurium

*Salmonella typhimurium* bet hedges by engaging in antibiotic persistence, where some proportion of cells adopt the T1-phenotype, which is a specialist that is fast-growing but sensitive to antibiotics, and the remainder adopt the T1+ phenotype, which is a generalist that is slow-growing but antibiotic tolerant ([40]). Arnoldini et al 2014 used single cell microfluidics to measure the proportion of cells that adopt the T1-phenotype (*P*_*Spec*_), as well as the doubling times for both cell types and their survivorship given clinically relevant antibiotic concentration (*w*_*A*_, *w*_*B*_, and *w*_*C*_). Interestingly, the observed *P*_*Spec*_ = 0.19 of *S. typhimurium* is higher than other estimates of persister fractions in *E. coli*, which normally range from as low as 0.001% to just over 10%.

Estimates of the probability of not encountering antibiotics (or *P*_*A*_) are challenging to estimate. Antibiotic adherence has been shown to vary with age, type or duration of treatment, socioeconomic status, and personality ([46], [47]). Estimates of the prevalence of non-adherence range from 30% to 70% of patients ([48], [49]). Within the large proportion of the population that are non-adherent, further measurements of the degree of non-adherence per individual can also vary. One study defined “excellent adherence” as taking only greater than 80% of prescribed doses; a benchmark only 30% of patients met ([49]). As such, it is likely that individual adherence can vary from *P*_*NoAntibiotics*_ = 0 to *P*_*NoAntibiotics*_ = 1.

We calculate the deterministic Δ*GMF* of the bet hedging *S. typhimurium* compared to a wild-type with a persister fraction of 0 across all values of *P*_*Noantibiotics*_ (fig 4C). We find that bet hedging is expected to be deleterious (Δ*GMF <* 0) when *P*_*Noantibiotics*_ *>* 0.932. In other words, when someone almost never takes antibiotics, then antibiotic persistence is not necessary for the infection to be successful.

Within our stochastic framework, we find that *S. typhimurium* does exhibit sign inversion when 0.9 *< P*_*Noantibiotics*_ *<* 0.95 (fig 4D). Given the large effective population size of bacteria, this means that *S. typhimurium* is expected to be beneficial for a larger range of parameter space than the GMF predicts. Bet hedging is less beneficial than the GMF predicts when *P*_*Noantibiotics*_ = 0.931 for *N <* 30. On the other hand, bet hedging is more beneficial than the GMF predicts when *P*_*Noantibiotics*_ = 0.94 when *N >* 100. Our results suggest that bet hedging is expected to evolve even when the probability of encountering antibiotics is very rare.

## Discussion

We expand on previous findings that deterministic treatments (namely, predictions based on the GMF) do not always adequately capture the sign of selection for bet hedging, a strategy for reducing risk in the face of unpredictable environmental change ([25], [29], [6], [30], [31], [32], [33]). It has been established that phenotypic stochasticity can elevate the extinction risk for bet hedging, while environmental stochasticity can lower the extinction risk ([33]). Additionally, population size may play an out-sized role in impacting the sign of selection, in addition to the strength of selection ([1], [29], [33]). Using both stochastic individual-based simulations and Markov numeric matrices, we find that explicitly incorporating stochasticity changes the sign of selection of bet hedging across a wide range of parameter space. When bet hedging is expected to be beneficial given its GMF, instead bet hedging can be deleterious at sufficiently small population sizes due to phenotypic stochasticity. In contrast, when bet hedging is expected to be deleterious given its GMF, bet hedging can actually be beneficial at sufficiently large population sizes due to environmental stochasticity. This non-monotonic dependence of the sign of selection on population size is known as sign inversion ([34]). Our results are generalizable for all types of bet hedging strategies: conservative, “high risk / low risk” diversified, and “high risk / high risk” diversified.

Our results provide another framework for evaluating the adaptive significance of real-world bet hedging traits beyond GMF, which we use on two study systems: variance in germination phenology in *Papaver dubium* and antibiotic persistence in *Salmonella typhimurium*. In both cases, we find that sign inversion occurs for realistic parameter estimates, and that bet hedging is expected to be beneficial for a larger range of environmental regimes than GMF predicts. For *P. dubium*, because historical climate records give insight into the environmental context in which this bet hedging strategy evolved, we are able to show that GMF predicts the observed degree of bet hedging to be deleterious, while our model shows bet hedging is beneficial.

Given that the largest existing problem in the field of bet hedging research has been discerning the adaptive significance of putative bet hedging traits, our model provides another tool in predicting when bet hedging is adaptive. In 2011, Simons found that only 10% of bet hedging studies were able to demonstrate that the putative bet hedging trait was able to increase geometric mean fitness in its observed environment ([5]). Approximately two thirds of the studies that failed to prove the adaptive significance of the bet hedging trait were able to empirically measure life-history parameters (such as *P*_*Spec*_), but fail to measure the environmental context in which the traits evolved, as was the case in our *Papaver* example ([14]). While not every example bet hedging trait in this category has readily available environmental data, our model provides an exciting opportunity to expand the number of examples of adaptive bet hedging. Further, our results show that sign inversion is most likely to occur when the absolute value of Δ*GMF* is small. As the *GMF* of the wild-type in our model is 1, Δ*GMF* can be considered equivalent to the selection coefficient in more classic population genetics models. Since the majority of selection coefficients are also near 0 ([50]), it seems likely that many real-world bet hedging traits will have Δ*GMF* values in the region of parameter space where sign inversion occurs. Study systems that have preliminary parameter estimates that could be expanded on through additional experimentation include within-clutch yolk size variation in ([51]), fruit set in Yucca ([52]), and diapuse in crickets ([53]). Future empirical work that expands on these data to connect fitness to environment could provide additional examples of cases where candidate bet hedging traits exhibit sign inversion.

Additionally, our finding that environmental stochasticity causes deterministic predictions to break down has broad implications for evolution in unpredictable environments. Bet hedging is often discussed in comparison to two other strategies for coping with unpredictable environments: adaptive tracking and phenotypic plasticity ([5]). Adaptive tracking is evolution by selection on standing genetic variation, and is predicted to be only adaptive when the time-scale of environmental change is slower than the time it takes for a mutation to reach fixation ([21], [5]). In other words, adaptive tracking is beneficial only when environments are essentially constant. Additionally, adaptive tracking does not result in phenotypic heterogeneity within a single genotype. Therefore, we predict that neither environmental nor phenotypic stochasticity will alter the evolution of adaptive tracking.

On the other hand, phenotypic plasticity, or change in an organism’s phenotype in response to some environmental cue, is more similar to bet hedging([7]). Phenotypic plasticity is predicted to evolve when environmental change is rapid and provides predictable cues; the organism must be able to change in response to environmental stimuli [5]. Similar to bet hedging, phenotypic plasticity is expected to evolve in fast changing environments; unlike bet hedging, plasticity is expected to evolve only when environmental cues are reliable. It follows that the evolution of phenotypic plasticity may be impacted by environmental stochasticity. Additionally, phenotypic plasticity and bet hedging may not be mutually exclusive. For example, phenotypic plasticity reaction norms can be “noisy”, resulting in phenotypic heterogeneity within a single generation ([5], [54]), [55]). In other words, “imperfect” phenotypic plasticity, or non-uniformity in response to a cue, can lead to a lineage exhibiting a combination of diversified bet hedging and plasticity. This means that phenotypic stochasticity may also impact the evolution of phenotypic plasticity. Therefore, it seems likely that phenotypic plasticity may similarly exhibit sign inversion. Future work that investigates whether phentoypic plasticity also exhibits sign inversion is needed.

Our results also fit into a larger conversation about traits that exhibit sign inversion. To date, all examples of sign inversion have been observed in lineage-variable traits, or alleles whose fitness can change between carriers ([21], [35]). In contrast to directly selected mutations, i.e. alleles with a constant fitness across all contexts, lineage-variable traits will experience variance in fitness either within a single lineage, such as alleles that increase cooperation ([56]) and recombination ([57], or between lineages, such as alleles that increase mutation rate ([34]). Bet hedging is unique among other lineage-variable traits because it displays both within-lineage fitness variance (i.e. variance in realized phenotype) and between-lineage fitness variance (i.e. variance in realized environmental regime) simultaneously. Our work shows that these two types of lineage-variable selection may not be mutually exclusive, and provides some preliminary insights into how within- and between-lineage variance can interact.

Our generalized model is able to predict the adaptive significance of bet hedging. Determining whether a bet hedging trait is adaptive in its current environmental context can be challenging, both from an experimental and theoretical perspective. Here, we show that GMF alone does not capture the full extent to which bet hedging is adaptive for all types of bet hedging strategies, and that stochastic models are necessary to explain the evolution of real-world bet hedging strategies.

---

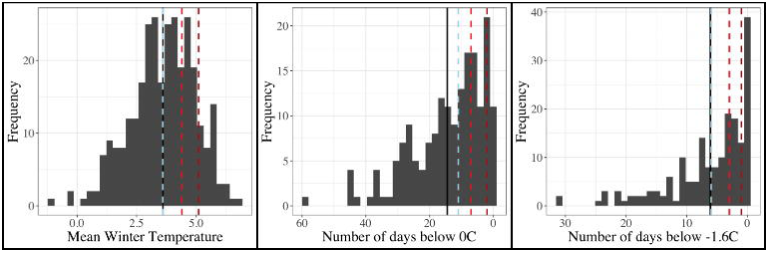

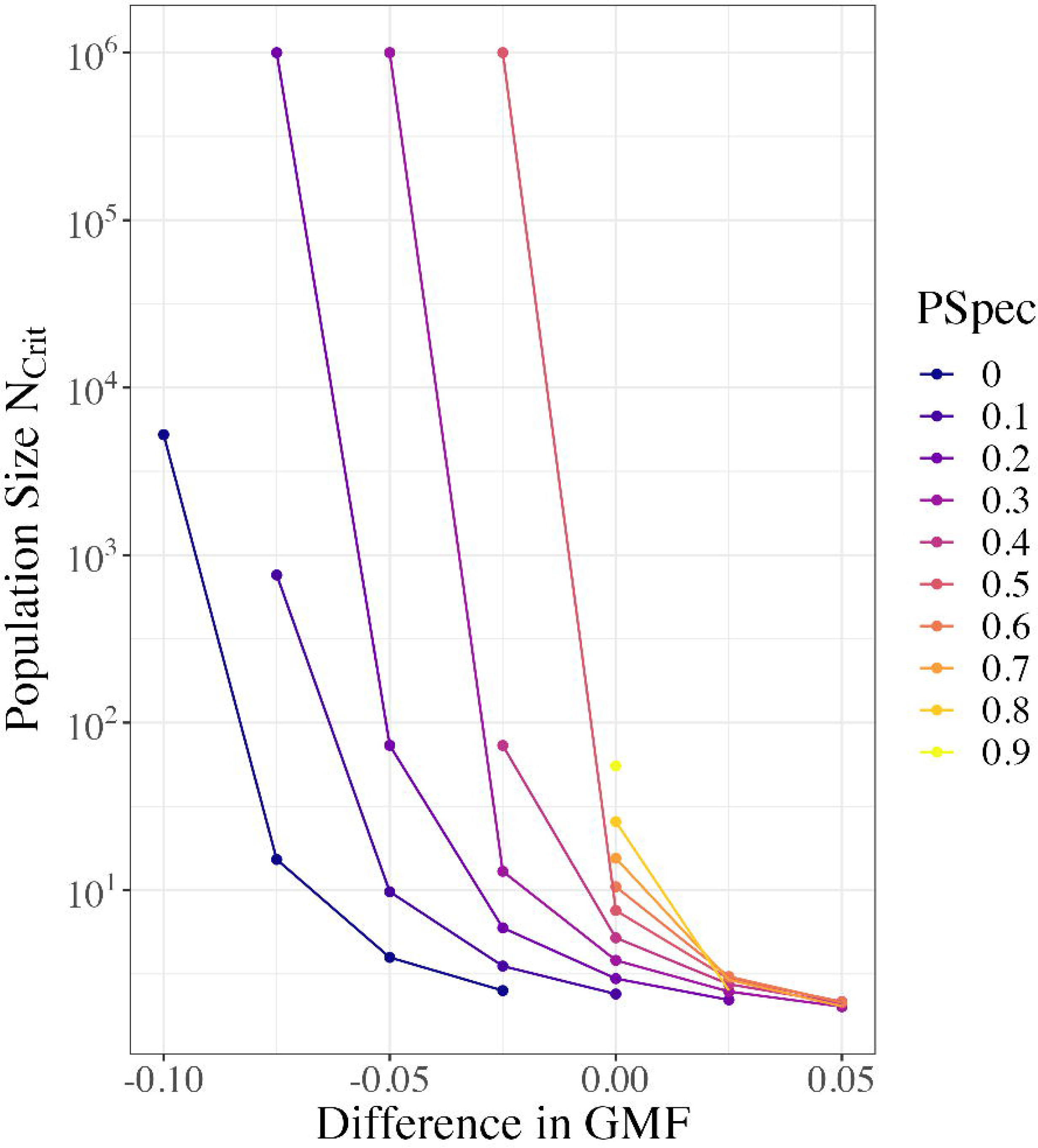

