## Supplemental Materials for "Beyond the (geometric) mean: stochastic models undermine deterministic predictions of bet hedger evolution"

### Supplemental Methods

#### Constructing Markov transition matrices

Markov transition matrices give the probability of transitioning from one state to another. In our case, the states are the number of bet hedger individuals in the population, which can range from $[0, N]$. Therefore, the matrices are $N+1 \times N+1$ matrices that give the probability of transitioning from all possible values of starting count, $x_t$, to all possible values of ending count $x_{t+1}$ in a single time step. Each column of the matrix corresponds to a different starting count of $x_t$, while each row corresponds to a different ending count of $x_{t+1}$. The matrix can therefore be written out as:

$$T = \begin{bmatrix} P(x_t = 0|x_{t+1} = 0) & P(x_t = 1|x_{t+1} = 0) & \cdots & P(x_t = N|x_{t+1} = 0) \\ P(x_t = 0|x_{t+1} = 1) & P(x_t = 1|x_{t+1} = 1) & \cdots & P(x_t = N|x_{t+1} = 1) \\ \cdots & \cdots & \ddots & \cdots \\ P(x_t = 0|x_{t+1} = N) & P(x_t = 1|x_{t+1} = N) & \cdots & P(x_t = N|x_{t+1} = N) \end{bmatrix}$$

Transition probabilities are calculated by constructing a binomial distribution, for which the probability of success is the expected frequency of the invading mutant given its current frequency and fitness, and the number of draws is equal to $N$.

### Supplemental Results

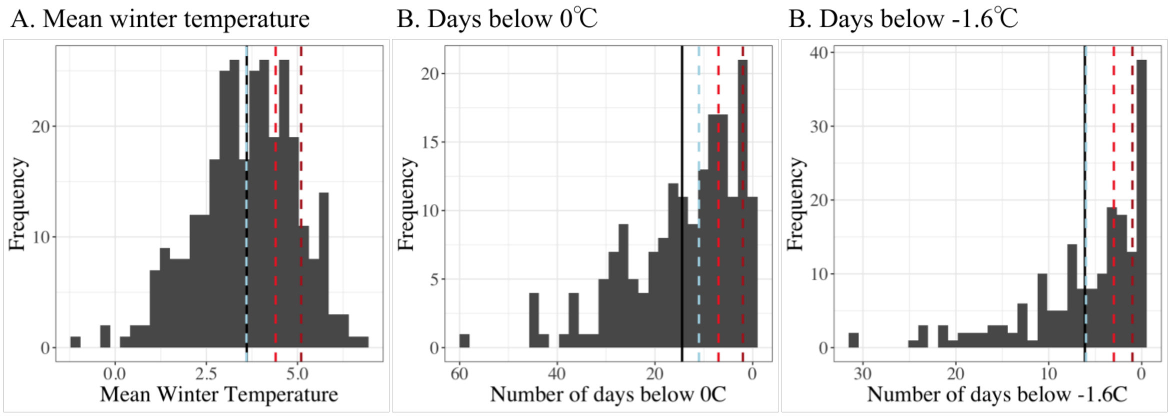


**Supplemental Figure 1:** Historical climate observations of daily winter temperatures were used to estimate the probability of a harsh winter. The black vertical lines correspond to the mean of the distribution¿ The blue vertical lines correspond to the year 1966, which was the observed ”harsh” year in Arthur 1973. The two red vertical lines correspond to the observed ”mild” years in Arthur 1973, 1966 (light red) and 1967 (dark red). A. Mean winter temperature. B. Number of days below 0°C per winter. C. Number of days below -1.6°C per winter.

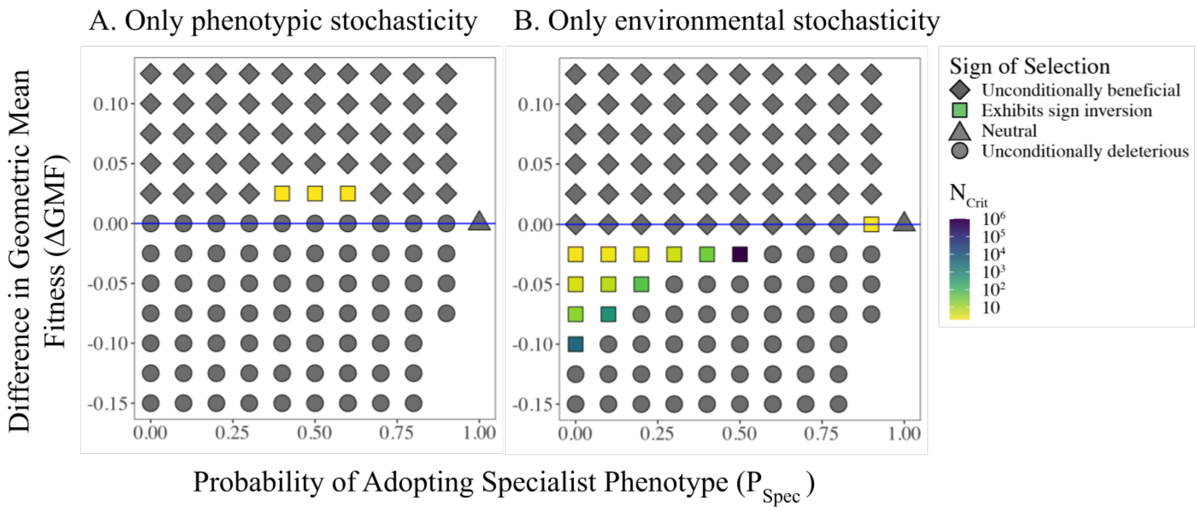


**Supplemental Figure 2:** The sign of selection of the bet hedger across parameter space is characterized using stochastic, individual based simulations. The x-axis is $P_{Spec}$, or the proportion of bet hedger offspring which adopt the specialist phenotype. The y-axis is the difference in geometric mean fitness ($\Delta GMF$) of the bet hedger and the wild-type. Values of $\Delta GMF$ that are impossible to achieve in combination with their associated $P_{Spec}$ are intentionally left blank (see Methods). For points in parameter space where bet hedging exhibits sign inversion, $N_{Crit}$ is interpolated. We utilize the "high risk / low risk" model of bet hedging. In both cases, stochasticity in reproduction is maintained. A. When phenotypic stochasticity is turned "on" and environmental stochasticity is turned "off", bet hedging exhibits sign inversion only when $\Delta GMF > 0$. Additionally, bet hedging is unconditionally deleterious when $\Delta GMF = 0$. B. When environmental stochasticity is turned "on" and phenotypic stochasticity is turned "off", bet hedging exhibits sign inversion only when $\Delta GMF < 0$. Bet hedging becomes unconditionally beneficial when $\Delta GMF = 0$.

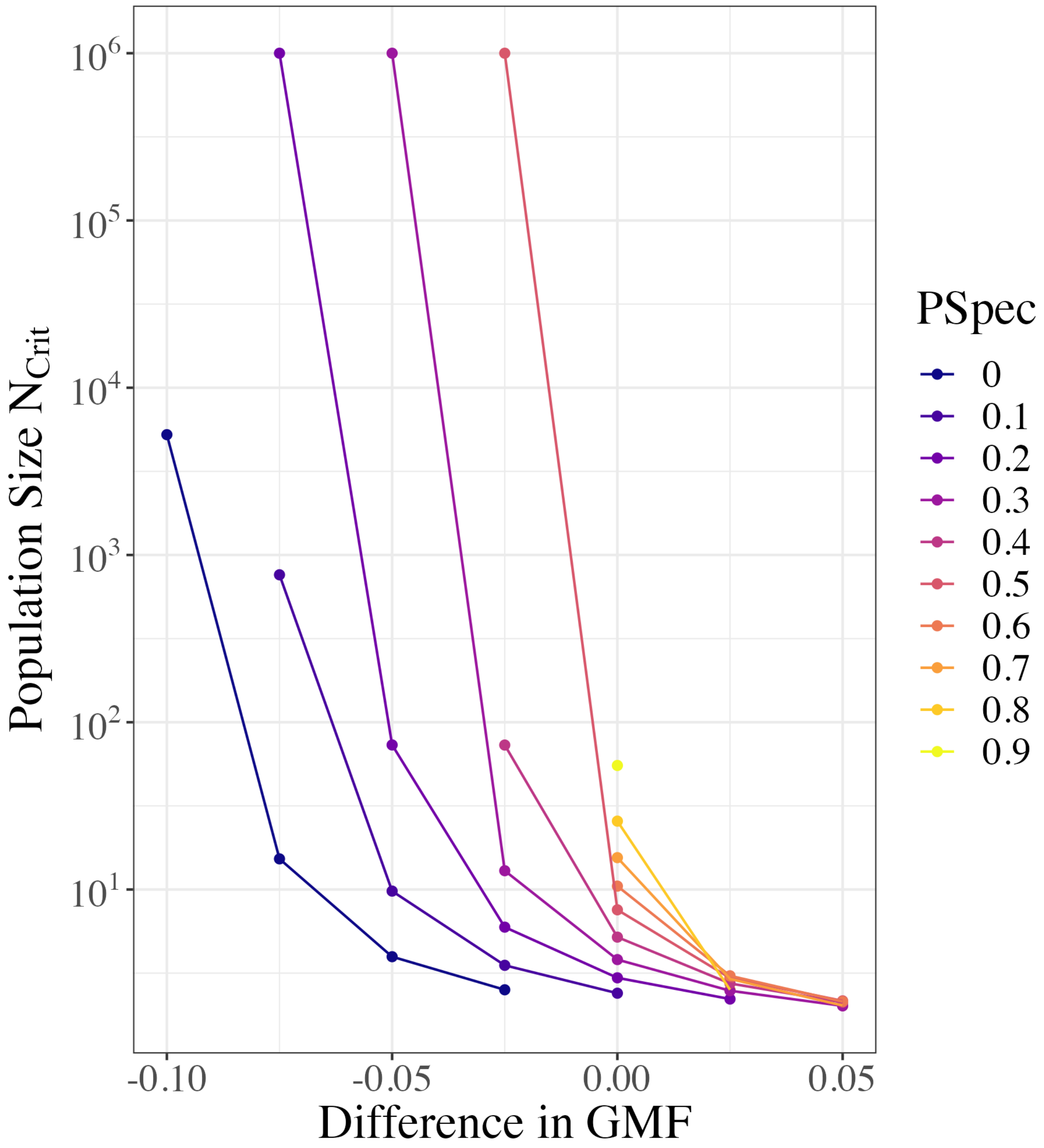


**Supplemental Figure 3:** The population size where the $NP_{Fix}$ curve crosses the neutral benchmarks ($N_{Crit}$) as a function of $\Delta GMF$ and $P_{Spec}$. As $\Delta GMF$ decreases, the value of $N_{Crit}$ increases exponentially. Keeping $\Delta GMF$ constant, as $P_{Spec}$ increases, the value of $N_{Crit}$ also increases.
